# Temperature-dependent Fano resonance in Microtubules and Tubulin investigated with Spontaneous Raman Spectroscopy

**DOI:** 10.64898/2026.11.04.764180

**Authors:** Mario D’Acunto

## Abstract

Microtubules are self-assembled biological hollow nanotubes composed by protein tubulin. They are essential for many biological functions, including cell motility, cell division, and act as intracellular track for organelle transport. In turn, through dynamic assembly and disassembly, they form spindle fibers for chromosome segregation during mitosis and meiosis. In this paper, we have measured the Raman spectra of microtubules and their constituent protein tubulin in dry powder form for a range of temperatures, from nitrogen liquid *T*=77 K up to room and physiological temperature *T*=310 K, to determine if molecular bond vibrations present potential Fano resonance asymmetry. We have found nontrivial persistence of Fano resonances, *q*<5, at room and physiological temperatures for some specific vibrational modes.

## I. INTRODUCTION

Fano-resonance line-shape arises from interference between a discrete states coupled to a continuum, so that its related asymmetry *q*-value is the resultant of interference between two scattering amplitudes, one due to the scattering within a continuum of states (the background process) and the second one due to an excitation of a discrete state, (the resonant state) [1-10]. This phenomenon, originally introduced in atomic physics [1-2], has been observed in a wide range of systems [3-9], including condensed matter, nanostructures, molecular spectroscopy, and biomolecular systems such as proteins [10], and amino acids [11].

The emergence of Fano-type line shapes in biological Raman spectra is generally interpreted as arising from interference between discrete vibrational states (molecular phonons of peptide bonds or side chains) and broad electronic or excitonic continua (arising from aromatic π-electron systems and their coupling to the protein environment). This interference occurs because the scattering amplitude of the continuum typically varies slowly with energy while the scattering amplitude of the discrete state changes quickly both in magnitude and phase, resulting in the characteristic asymmetric profile. This is the general framework of Raman spectroscopy, where a sample is irradiated with monochromatic light, resulting in the initial discrete vibrational ground state being coupled to a final discrete vibrational excited state through an intermediate state in the continuum of electronic excited state. This general mechanism of Fano resonance persists even if the incident radiation does not have enough energy to actually reach one of the electronically excited states, as in spontaneous Raman spectroscopy, because the initial and final vibrational states are coupled through an intermediate virtual statev that is part of continuum of electronic states, Fig. 1.

**FIG 1.**
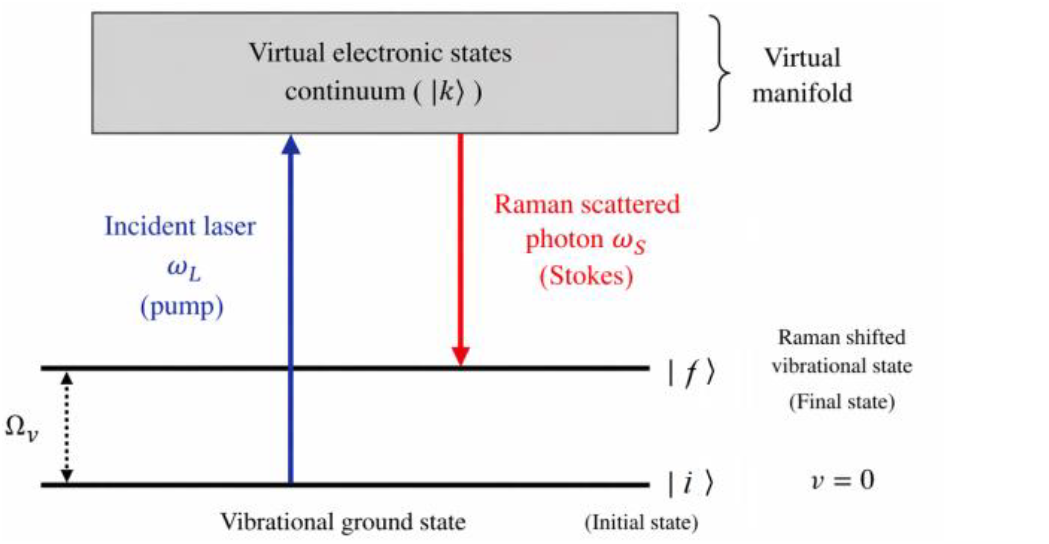
Schematic energy-level diagram for Raman spectroscopy modelled as a Fano-type system. The molecular vibrational ground state ∣*i*⟩ and Raman shifted state ∣*f*⟩ are coupled through a continuum of virtual electronic intermediate states (gray block). The incident laser field of frequency *ω*_*L*_ excites the system to the virtual manifold, followed by inelastic scattering at frequency *ω*_*S*_. For Stokes Raman scattering, considered in this investigation, *ω*_*S*_=*ω*_*L*_-Ω_*v*_, where Ω_*v*_ is the vibrational frequency. For details see Fano-type Hamiltonian in Appendix.

This mechanism is formally identical to Fano interference originally described in atomic physics and later generalized across condensed matter systems, where asymmetric spectral profiles emerge from quantum interference between resonant and non-resonant scattering channels.

Asymmetric Raman line shapes attributed to Fano interference are not unique to biological systems. Similar effects have been reported in metallic carbon nanotubes, where phonon–electron coupling produces asymmetric G-band features [12]; nanophotonic and metamaterial systems, where interference between discrete and continuum optical modes generates tunable Fano resonances [13]; photonic crystals and plasmonic structures, where engineered symmetry breaking produces robust asymmetric spectral profiles [14]. The recurrence of Fano-like line shapes across such diverse systems supports the generality of the underlying interference mechanism, although biological interpretation requires caution due to additional structural and thermal complexity.

In Raman spectroscopy, Figure 1, the Raman intensity as a function of the Raman shift *ω* can be described by the standard Fano line-shape formula

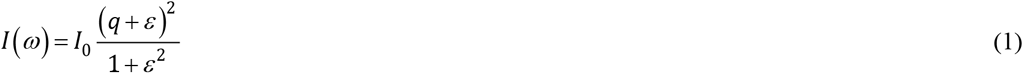

Where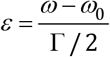 and *ω*_0_ denotes the resonance position, *Γ* is connected to the full width at half maximum (FWHM) of the resonance, *q* is the Fano asymmetry parameter, *I*_*0*_ is an overall intensity scale, and *B* represents a constant background contribution.

The parameter *q* quantifies the relative strength and phase between the discrete and continuum scattering amplitudes. In the limit |*q*| → ∞, the interference vanishes and Eq. (1) reduces to a symmetric Lorentzian profile. Finite values of *q* indicate nontrivial discrete–continuum coupling, while the linewidth Γ encodes both intrinsic vibrational lifetimes and environment-induced dephasing processes.

In spontaneous Raman scattering, the intensity of a Stokes line, *I*, associated with a vibrational mode of frequency *ω*_0_ is governed by the modulation of the electronic polarizability induced by nuclear motion. Within the electrically non-resonant regime, the temperature-dependent Stokes Raman intensity can be written as [15-16]

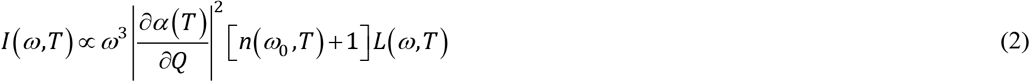

where *ω* is the emitted photon frequency, *Q* is the normal coordinate of the vibrational mode, *α* is the molecular or lattice polarizability, and 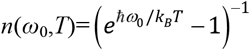 is the Bose–Einstein occupation factor and 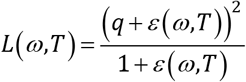 . The cubic dependence on *ω* originates from the photon density of states in spontaneous emission.

Eq. (2) shows that the temperature dependence of the Stokes Raman intensity arises from three distinct contributions: (i) the Bose occupation factor *n*(*ω*_0_,*T*)+1, (ii) the temperature evolution of the vibrational frequency and linewidth entering *ε*(*ω*(*T*),*T*), and (iii) the polarizability derivative ∂α(*T*)/∂Q, which encodes changes in the electronic structure and lattice anharmonicity. While the Bose factor can be removed explicitly by normalizing the measured intensity by *n*(*ω*_0_,*T*)+1, the residual temperature dependence of the normalized intensity provides direct experimental access to variations of |∂*α*/∂*Q*|^2^.

Finally, it is worth noting that the Fano parameter *q*(*T*) itself depends on the ratio between the discrete vibrational scattering amplitude and the continuum contribution. Since the discrete channel is proportional to the polarizability derivative, temperature-induced changes in ∂α(*T*)/∂Q may manifest not only as an overall intensity variation, but also as a systematic evolution of the Fano asymmetry.

Microtubules (MTs) are self-assembled biological structures like rigid hollow nanotubes, with approximately 25nm in diameter, composed by αβ-Tubulins (TBs), Figure 2 [17]. Like actin filaments, MTs are dynamic structures that experiment continual assembly and disassembly within the cell. They are essential to determine cell shape and for a variety of crucial cell functions, including cell movements, cell locomotion, intracellular transport of organelles, and, in turn, separation of chromosomes during mitosis. αβ-TBs are the proteins that make up microtubules and belong to tubulin superfamily that includes five distinct families of globular proteins, of which αβ-TBs are the most common constituents. They have a molecular weight of approximately 55kDa and slightly differ one each other. α-tubulins present a total of 439 residues, which most abundant is Alanine (8%), second most abundant is Glycine (7.7%) and least abundant is Tryptophan (0.9%). β-tubulins have 427 residues, where most abundant amino acid is Leucine (8%), second most abundant is Valine (7.7%), and Tryptophan as the least abundant (0.9% in α-TB and 0.7% in β-TB, respectively). Other difference are represented by the relative abundance of Lysine and Proline in α-TBs, and Methionine and Phenylalanine in β-TBs. Both have a very similar overall composition, which is expected as they are homologous proteins. The slight differences evidenced reflect their distinct functional roles and sequence variations. To form microtubules, the dimers of αβ-tubulins bind to Guanosine-5′-triphosphate (GTP) and assemble onto the (+) ends of microtubules, while in the GTP-bound state, so that α- and β-tubulins subunit are exposed in opposite ends of microtubules, respectively.

**FIG 2.**
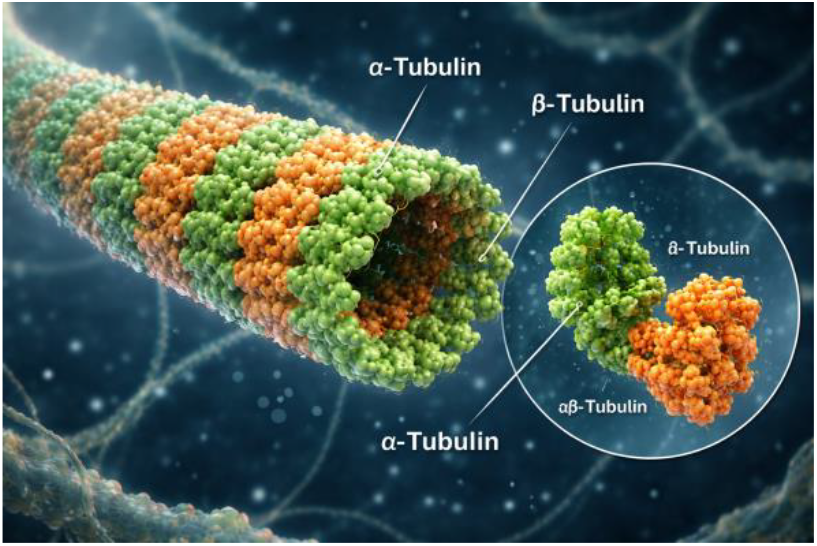
Schematic representation of a microtubule highlighting the organization of α- and β-TB subunits. Microtubules are cylindrical polymers composed of repeating αβ-TB heterodimers arranged longitudinally into protofilaments, which associate laterally to form a hollow tube. In the image, α-TB is shown in green and β-TB in orange. The inset provides a magnified view of a single αβ-TB dimer, emphasizing the structural pairing of the two subunits. Credits: the image was generated with openAI.

Recent theoretical works suggested that MTs could present unique optical properties mediated by interactions between aromatic amino acid tryptophan, which would lead to exceptional properties of excitation transfer and superradiance, and finally to possible metamaterial behavior [18-21].

In addition, MTs provide a structurally periodic, quasi-crystalline environment that may enhance vibrational coupling effects relative to monomeric TB. Their cylindrical geometry and ordered protofilament arrangement introduce spatial constraints that can modify vibrational density of states and promote partial coherence among vibrational modes [21]. This structural ordering has led to theoretical proposals that MTs could behave as coupled vibrational-electronic systems with enhanced optical response, sometimes described in terms of “biological nanowire-like” behavior [19]. Within this context, Raman asymmetry has been interpreted as a spectroscopic signature of enhanced coupling between molecular vibrations and delocalized electronic excitations in the protein lattice.

The results of the present investigation show that Raman spectra of TB and MTs contain reproducible Fano asymmetric line shapes of some vibrational modes. One of such vibrational mode ∼833cm^-1^, assigned to tyrosine doublet and very sensitive to H-bonds is common to both the two biomolecular samples. In addition, while TBs show Fano symmetry at physiological temperature for ∼1046cm^-1^ mode, correspondent to Phe-H in plane vibrations and C-C stretching in aromatic amino acids (see Table I), MTs present analogous *q* values for ∼715cm^-1^ mode, correspondent to C-S stretching in sulfur class amino acids.

The persistence of finite *q* values up to physiological temperatures implies that the ratio between coherent resonant Raman scattering to dissipative continuum scattering remains non-negligible, supporting the view that MTs and TBs behave as structured open vibrational systems exploiting environment interaction rather than overdamped biological media.

## II. METHODS

Solid powder TBs and MTs, stored at room temperature, were provided by Cytoskeleton (Denver, CO, catalog numbers T240 and MT002, respectively). Since MT002 (>90% pure) is taxol-, sucrose- and dextran-stabilized, possible contribution of such excipients was carefully checked by recording Raman spectra of taxol (∼4% overall content) and sucrose (∼5% overall content ) and comparing with MT spectra. Both taxol and sucrose were purchased by Merck. Raman spectra were collected with a confocal Raman microspectroscopy (Xplora Plus, Horiba Scientific) equipped with a 532nm diode laser source and 2400 lines/mm grating giving a spectral resolution of ∼1.15 cm^-1^. Potential contributions to asymmetry from the experimental apparatus were carefully checked. The optimal experimental setup has been configure with slit and pinhole at 100μm, respectively.

The temperature was changed in the range *T*=77 K (−196°C) to *T*=310 K (37°C) making use of a Linkam HFS600E-PB4 Probe Stage to modulate temperature and powder samples holder during all the measurements. To work with Linkam probe Stage, the confocal Raman microspectroscopy was equipped with a long focal ×50 air objective (0.5NA, Olympus) with a nominal power laser of ∼30mW. Acquisition time and repetition to collect Raman signal were designed to 1 sec and 40, respectively for all the samples. Four replications of each measurement -starting form initial physiological temperature (T=310 K) and and progressively reducing the temperature until liquid nitrogen (T=77 K) and then do the reverse process increasing the temperature in order to highlight hysteresis effects-were made and averages of them were taken to find a nominal standard deviations.

### A. Spectral fitting

The whole Raman spectra could be expressed as a sum of multiple Fano functions, as given by Eq.(2), so that

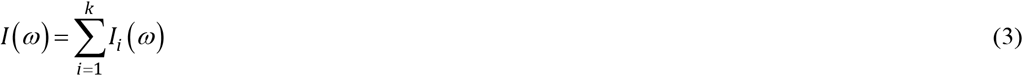

where *k* is the total number of peaks considered in the fitting model.

Spectral data were processed and analyzed using a custom automated routine in MATLAB to extract peak parameters and evaluate line-shape asymmetry. The analysis followed a four-stage pipeline:

Raw spectra were first truncated to a region of interest between 400 and 1800 cm^−1^. Although raw data presented low background, except aromatic class amino acids, to improve the signal-to-noise ratio (SNR) for robust peak identification, the data underwent a baseline subtraction (minimum-intensity offset) and were smoothed using a Savitzky-Golay filter with a third-order polynomial and an 11-point frame length. Initial peak locations were identified using a local maxima detection algorithm. Candidates were filtered based on a minimum peak height and prominence threshold. For each identified peak, a local computational window was defined, extending ±3 standard widths from the peak center to isolate the relevant spectral feature for fitting. Since baseline subtraction can introduce artifacts, which can exaggerate apparent asymmetry, the quantification were made both on raw and baseline-subtracted spectra, evidencing artifacts on these last one, so the results on *q*-values here presented regard exclusively raw data.

To account for potential asymmetric profiles—often indicative of quantum interference or phonon-electron coupling— each localized band was modeled using the Fano resonance Eq. (6). Parameters were optimized via a Non-linear Least-Squares (*Lsqnonlin*) approach using the Levenberg-Marquardt algorithm. Initial guesses for the fit were derived from the smoothed data characteristics, in addition, fit parameters were constrained within physically realistic bounds to prevent divergence. For each fit, the Full Width at Half Maximum (FWHM) was calculated as:

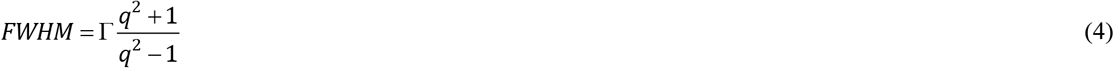

The goodness-of-fit was quantified using the Root Mean Square Error (RMSE) and the Coefficient of Determination (*R*^2^). Only fits achieving high statistical significance were prioritized for final reporting, with results sorted by *R*^2^>0.9 to ensure the selection of the most physically relevant models, Figure 5. Using the spectral fitting technique, Raman spectra were fit by multiple models between a minimum and a maximum number of peaks identified. Each model was the assessed via the Alkaike information criterion (AIC) [22]. This procedure was performed for peaks within the range 410-1790cm^-1^ in order to avoid the potential edge effects in the fitting procedure. Since AIC values indicate a better fit model and by comparing the models among them with lower AIC values, the model with the optimal number of peaks giving the minimum AIC score was selected as the candidate for the best fit. In turn, we performed multi-peak fitting using only Lorentzian profiles for each identified band. AIC scores were calculated and compared. The Fano model consistently yielded significantly lower AIC values, confirming it is a statistically superior descriptor of the observed line shapes.

## III. RESULTS

Figure 3 shows the Raman spectra with the baseline removed for the TB (black line) and MT (red line) powders, respectively. at 123 K (the temperature is indicative). The peak identification and tentative assignation are reported in Table 1. Basic differences are evidenced by bold.

**Table 1.** Comparative peak assignation with in bold the bands evidencing Fano resonance asymmetry *q*-values ≤10 for a wide range of different temperatures. Ref. assignations [10, 23-28].

| Observed Peak (cm<sup>−1</sup>) | Tubulin | Microtubule | Tentative Assignment | Notes |
|---|---|---|---|---|
| 510–525 | weak | moderate | S–S stretch (disulfide) | Cys environment |
| **710–720** | strong | **very strong** | C–S stretch | **Strong change in MT** |
| **830–837** | strong | moderate | Tyr out-of-plane- C-H bending | **H-bond sensitive** |
| 930–945 | weak | broad | α-helix C–C stretch | secondary structure |
| 1001–1005 | moderate | strong | Phenylalanine ring breathing | protein marker |
| **1030–1048** | **weak** | **moderate** | **Phe C–H in-plane / C–C stretch** | **aromatic** |
| 1080–1100 | very strong | broad strong | C–N stretch / skeletal modes | peptide backbone |
| 1125–1150 | moderate | moderate | C–C / C–N stretch | ordered chains |
| 1205–1215 | weak | moderate | Phe/Tyr ring stretch | surface exposure |
| 1240–1270 | weak | broad | Amide III (β-sheet/random coil) | conformational |
| 1445–1465 | strong | stronger | CH<sub>2</sub> / CH<sub>3</sub> bending | protein packing |
| 1510–1525 | weak | moderate | Tyr / Trp ring mode | aromatic environment |
| 1600–1615 | weak | moderate | aromatic C=C stretch | Tyr/Phe/Trp |

**FIG 3.**
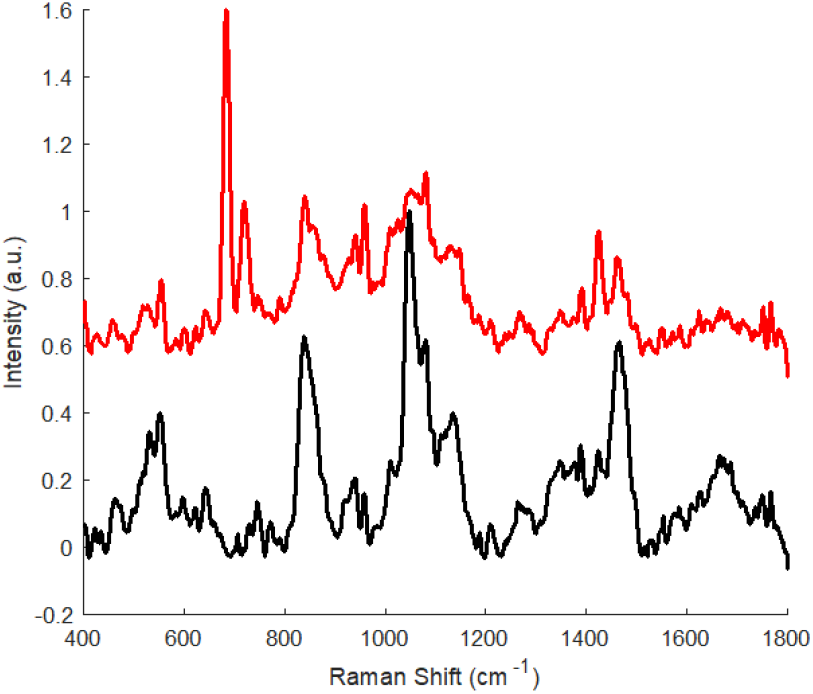
Raman spectra of tubulin (black line) and microtubule (red line) after baseline subtraction at T=273 K (the selected temperature is purely indicative). Main difference could be associated to polymerization processes in microtubules formation.

When TB assembles into MT, Raman changes usually reflect: i) increased β-sheet / ordered structure; ii) altered amide I/ amide III; iii) changed aromatic residue environment (Tyr, Trp, Phe); iv) altered Cys–S–S / C–S vibrations; v) tighter intermolecular packing. In part such expected differences are matched in Figure 2, which shows: i) stronger band near 700 cm^−1^ induced by C-S stretching; ii) broader/intense region 900–1150 cm^−1^; iii) stronger 1450 cm^−1^ ; iv) smoother / more ordered backbone signature.

Figures 4 and 5 report the normalized spectra at different temperatures and what we observe as expected, not reported in the figures, is a slight increasing of photon scattered at low temperature due to fewer thermally populated phonons and less vibrational disorder induced by phonon-phonon scattering.

**FIG 4.**
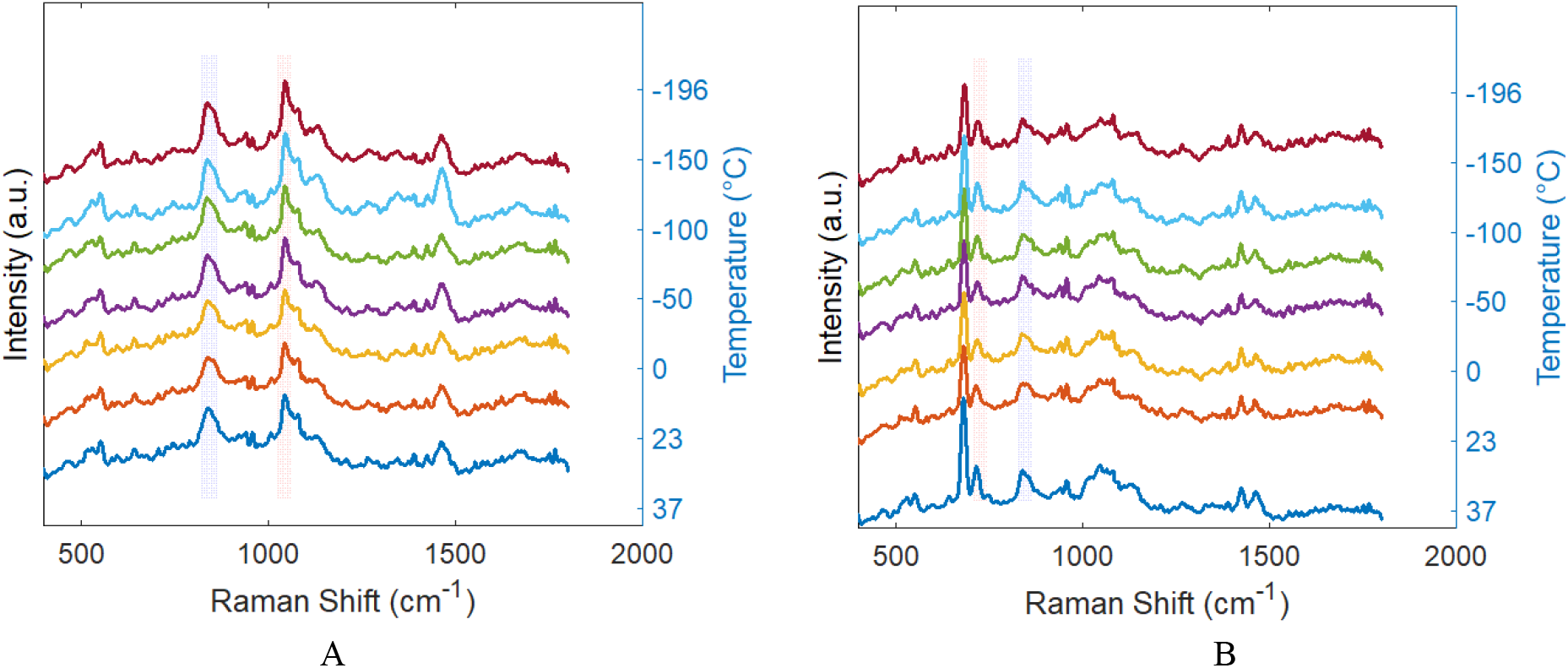
Normalized TB and MT Raman spectra at different temperatures, A and B, respectively. In both the figures, the rectangular shadow overlaps the peaks with Fano resonance ∣*q*∣≤10 for all temperatures.

**FIG 5.**
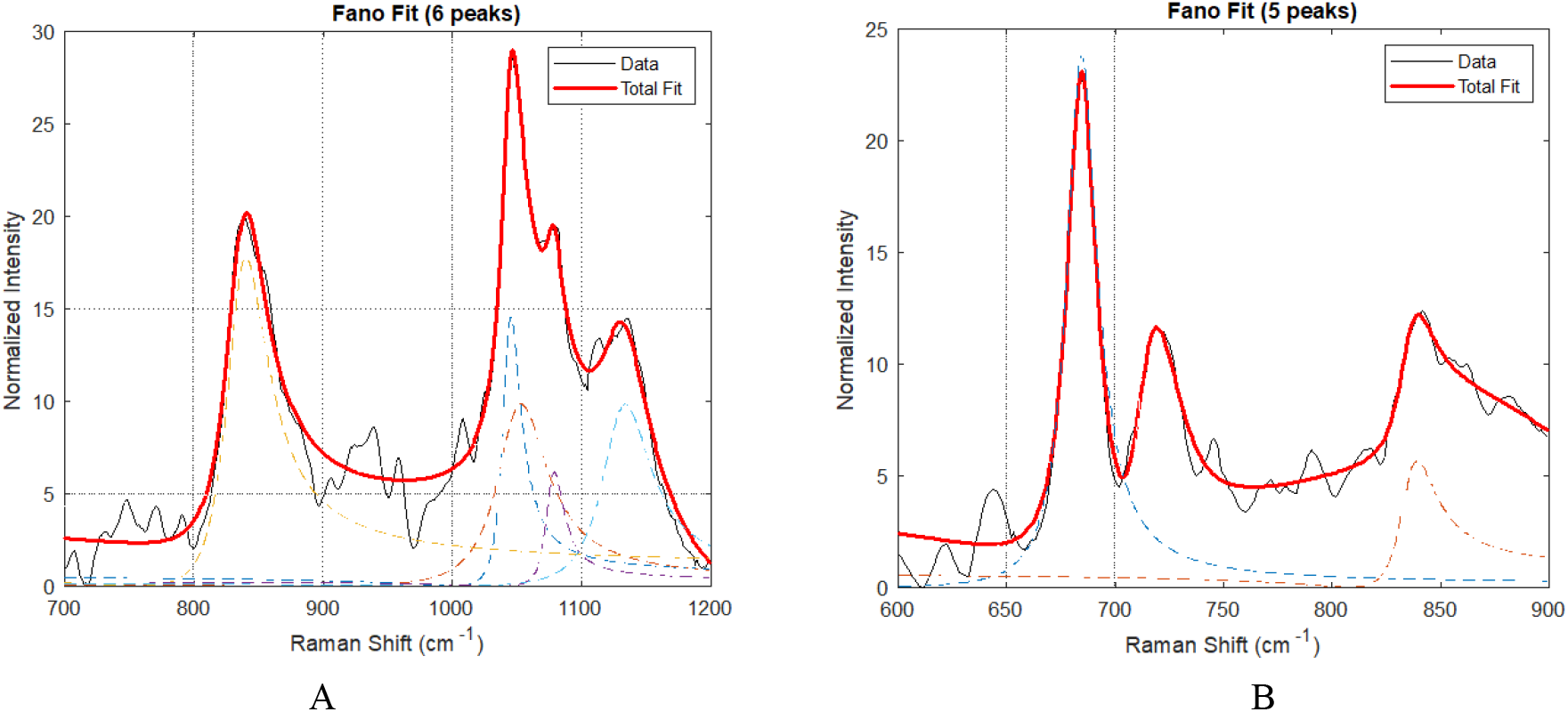
Decomposition of TB and MT Raman spectra on the range of interest for Fano resonances, A and B, respecyively.

**FIG 6.**
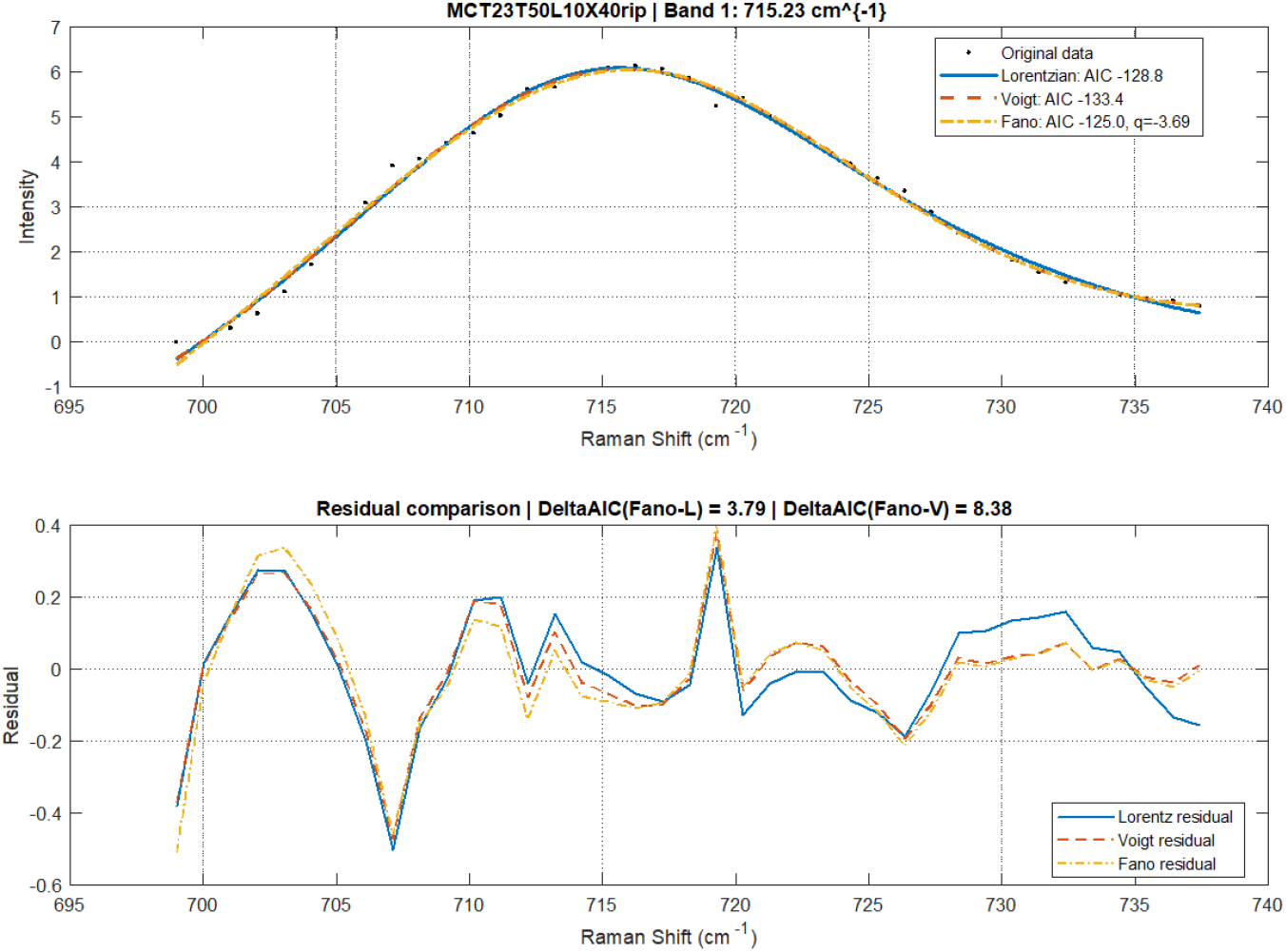
Example of discrimination of best fitting profile among Fano, Lorentzian and Voigt options via Matlab code written ad hoc. In this case we show the fitting of MT band at ∼715cm^-1^ at 300 K (23 °C).

Spectral fitting for the ranges of interest, i.e. reporting the most interesting *q* values for Fano resonance are reported in Figure 5. The red line represents the total fit.

The Fano resonance *q* values only for Raman bands with ∣*q*∣≤10 are reported in Figure 7. It should be noted that not-trivial behavior of temperature dependence of *q* values. Whereas, the correspondent Γ-values, see Eq. (4), are reported in Figure 8.

**FIG 7.**
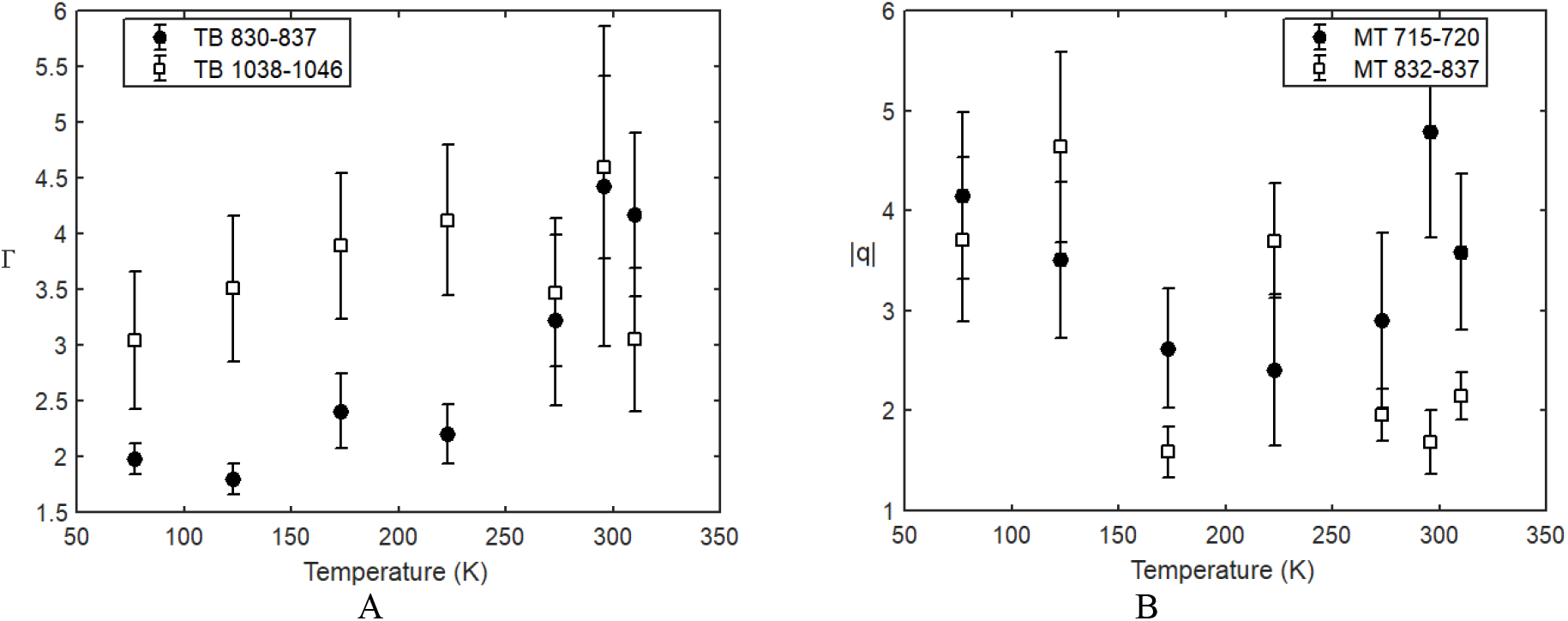
Raman active modes evidencing ∣*q*∣≤10 values at all temperatures. A: Raman bands at ∼830cm^-1^ and ∼1046cm^-1^for TB. B: Raman bands at ∼715cm^-1^ and ∼830cm^-1^ for MT.

**FIG 8.**
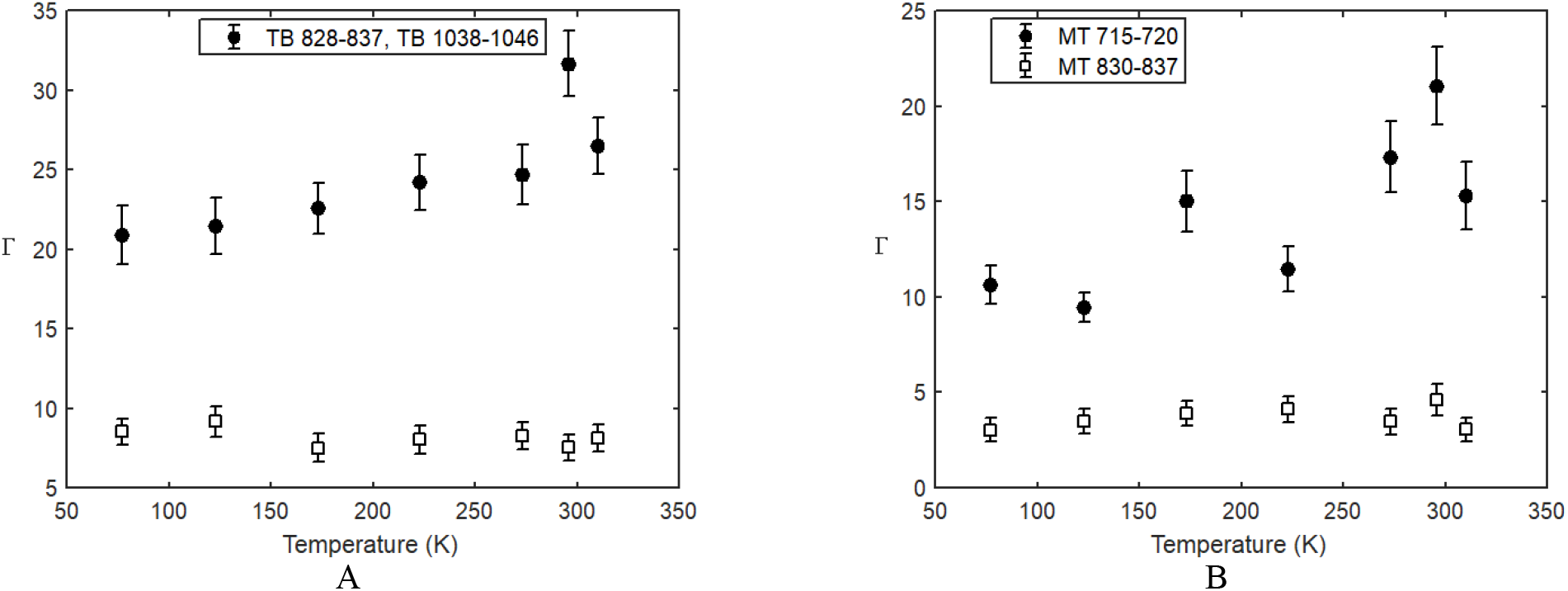
Linewidths Γ for the vibrational modes of interest displayed in FIG. 8, for TB (A) and MT (B), respectively.

## IV. DISCUSSION

The asymmetric Raman bands observed in TB dimers and polymerized MTs can be consistently interpreted within the open-system Fano Hamiltonian developed in the Appendix. In this framework, discrete protein vibrational excitations interact with a structured continuum of environmental states while simultaneously coupling to the optical Raman field. The experimentally extracted quantities Δ, Γ, and *q* thereof acquire direct microscopic meaning: Δ characterizes the thermal induced detuning of the effective vibrational eigenmode, Γ measures total relaxation and dephasing, and *q* quantifies the interference between resonant and continuum scattering channels. The experimental data do not show significant change of Δ as a function of temperature, so that our analysis can be focused on the changes of Γ and *q*.

An useful phenomenological relation follows from standard Fano theory, in which the asymmetry parameter scales as the ratio between the resonant discrete scattering amplitude and the continuum background amplitude. In the present system this may be written as [2, 29-30]

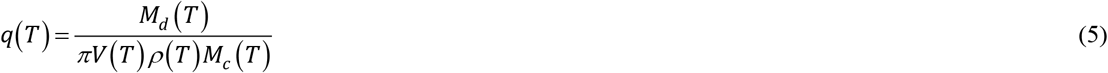

where *M*_*d*_(*T*) is the Raman matrix element for the discrete vibrational channel, *M*_*c*_(*T*) is the background continuum scattering amplitude, *V*(*T*) is the coupling between the discrete mode and the continuum, and *ρ*(*T*) is the effective continuum density of states. Eq. (5) is not intended as a full microscopic derivation, but as a compact guide for interpreting the measured temperature dependence of *q*, see Appendix.

Within this picture, the persistence of moderate asymmetry values *q*≅2–5 over the full interval *T*=77–310 K indicates that the ratio between coherent resonant scattering and dissipative background scattering remains finite even at physiological temperature. If thermal fluctuations completely destroyed pathway interference, the resonant contribution would become negligible compared with the continuum term and the spectra would evolve toward symmetric Lorentzian profiles. Instead, the observed finite values of *q* imply that the environmental continuum remains spectrally structured and that coherent pathway competition survives substantial thermal activation.

The extracted linewidths provide complementary information. In the model of the Appendix, the total width is determined by intrinsic vibrational decay, coupling-induced broadening, and pure dephasing, so that linewidth changes can be directly interpreted in terms of modified dissipation channels. For tubulin, the bands ∼1046 cm^−1^ and ∼830 cm^−1^ display widths of order 8–10 cm^−1^ and 20–30 cm^−1^, respectively, while in polymerized microtubules the ∼830 cm^−1^ feature narrows and an additional asymmetric mode emerges near ∼715 cm^−1^. Such behaviour suggests that polymerization redistributes vibrational decay pathways rather than simply increasing static disorder. By using the conversion factor 1cm^-1^ = 0.124meV, we obtain Γ in Eq. (4) correspond to 0.9-1.2meV for the mode at 1046cm^-1^ and 2.5-4meV for the mode around 830cm^-1^. Such values are quite lower than the thermal energy at physiological temperature ≅25meV, so that the ratio Γ/*k*_*B*_*T* is ∼0.04-0.14. Such ratios denote a weak phonon-bath coupling regime, despite the high thermal population environmental modes.

In term of the Fano-like Hamiltonian (see Appendix), the eigenvalues *λ*_±_ *=* −*γ* ± *i*Δ, where *γ* is the decay factor and Δ is detuning factor, immediately give Re(*λ*_±_)≈−(0.6−0.75)meV for the 1046cm^-1^ mode, and Re(*λ*_±_)≈−(1.2−1.6)meV for the 830cm^-1^ mode, denoting that the Liouvillian poles remain relatively close to the imaginary axis even at 310 K. In turn, microtubule polymerization changes the Liouvillian spectrum, while ∼830cm^-1^ mode is common to Tubulin or microtubules, but linewidths can differ due to polymerization, ∼715 cm^-1^ mode can be considered as the emergence of an additional slowly decaying Liouvillian mode associated with collective vibration dynamic of the polymerization lattice. MT modes show oscillatory behaviour, exhibiting simultaneous changes in *q* and suggesting that d*λ/*d*T* changes consequently, this behaviour could imply a reorganization of the underlying Liouvillian spectrum rather an isolated vibrational effect. At physiological temperature, the ratio between linewidths and thermal energy can be re-written as |Re(*λ*_±_)|/*k*_*B*_*T*≈0.05-0.08, so despite thermal energies exceeding the Liouvillian decay scales by approximately one order of magnitude, finite Fano asymmetries persist. This indicates that the relevant Liouvillian modes remain spectrally resolved sustaining coherently the Raman response.

As evidenced in Figure 8 the linewidths are not governed by a single relaxation channel, denoting coexisting of several mechanisms, which in part should follow the Fano interference changes with temperatures. Standard anharmonic phonon broadening models fail to reproduce the experimentally observed non-monotonic linewidth evolution. So that the anomalous linewidth fluctuations indicate that vibrational decoherence in TBs and MTs is dominated not by equilibrium phonon decay, but by heterogeneous conformational dynamics and correlated environmental fluctuations. For a Fano mode, changes in Γ may indicate variations in the interference conditions, for example, changes in coupling strength and changes in the continuum density of states, rather than simple vibrational damping.

One particularly important observation is that Figure 8 indicates a substantial linewidth reduction for the 832–837 cm^−1^ mode after microtubule polymerization. A natural interpretation is that self-assembly into the ordered cylindrical microtubule lattice renormalizes both *V*(*T*) and *ρ*(*T*). In isolated TB dimers, local configurational fluctuations and solvent exposure likely generate a broader and less selective continuum. After polymerization, protofilament alignment, lateral dimer–dimer contacts, and correlated hydration-shell dynamics may suppress some dissipative channels while enhancing others. This would reduce parts of Γ, modify the available density of states *ρ*, and thereby shift the balance entering the Fano parameter *q*. However, in light of the experimental data, we cannot go beyond this qualitative description, and this result would warrant a specific investigation, which is currently beyond the scope of the present work.

The appearance of additional asymmetric bands in MTs also supports a transition from predominantly local vibrations to partially collective excitations. In the Hamiltonian language, the bare term *H*_eff_ (see Appendix) is altered by intermolecular mechanical coupling, producing new normal modes distributed over multiple TB dimers. These collective coordinates may couple differently to the surrounding continuum than isolated molecular vibrations, explaining why polymerized microtubules exhibit distinct line-shape parameters from free tubulin.

Temperature dependence provides a second important control parameter. As temperature rises, bath occupation numbers increase and dephasing processes generally become stronger. One therefore expects an increase in Γ(*T*) and potentially stronger continuum scattering *M*_*c*_(*T*). However, the data indicate that asymmetry is not erased. This implies that thermal activation does not simply randomize the system; instead, it modifies all competing terms in a correlated way. For example, conformational softening could increase the discrete Raman matrix element *M*_*d*_(*T*), partially compensating stronger damping. Likewise, hydration dynamics may restructure the continuum density of states rather than merely broadening it.

These observations place TBs and MTs in an intermediate mesoscopic regime where coherence and dissipation coexist. This is distinct from highly ordered crystalline solids, where weak disorder often preserves sharp excitations, and also distinct from strongly overdamped soft matter, where vibrational structure is rapidly washed out. Cytoskeletal assemblies combine nanoscale order, biochemical heterogeneity, hydration-mediated dynamics, and self-organized architecture [31]. Such systems may therefore host interference effects unavailable in either conventional solids or simple liquids.

The present results also suggest a route toward functional spectroscopy of biological self-assembly. Because the parameter *q*(*T*) depends on the relative strength of discrete and continuum channels, it may serve as a sensitive reporter of structural transitions, ligand binding, nucleotide state, or mechanical strain in tubulin-based systems. Unlike peak position alone, which primarily tracks local force constants, the Fano asymmetry probes coupling topology and environmental mode structure.

At the same time, caution is required in interpretation. Fano-type profiles do not uniquely imply quantum coherence, since analogous asymmetric spectra can arise in classical coupled-oscillator networks or interference between damped response channels [32-34]. The significance of the present measurements is therefore not the automatic demonstration of long-lived quantum coherence, but rather the clear evidence that multiple scattering pathways interfere measurably in a complex biological material. Indeed, the Fano lineshape formula, Eq. (1), is mathematically identical whether derived from a quantum interference framework between discrete and continuum transition amplitudes or classical coupled oscillators with appropriate damping, so that can be truly difficult to evidence a general criterion to distinguish between quantum or classical effects in the experimental dataset on microtubule and tubulin, or amino acids [11]. One approach on this intriguing question could rely on quantify quantunmess of Fano-type Hamiltonian, as in appendix, through the negativity of Wigner function [35-37].

If we consider Eq (5) as a quantum derivation of *q*-values, some key temperature dependencies can be evidenced. Raman matrix element, *M*_*d*_(*T*) would show weak *T*-dependence, discrete-continuum coupling, *V*(*T*), can change sign with temperature, in turn, continuum density of states, *ρ*(*T*), and background scattering, *M*_*c*_(*T*), both increase with *T*. Given such behavior, *q*(*T*) is expected to decrease monotonically with increasing *T* with possible sign changes if *V*(*T*) crosses zero. In classical case, the correspondent *q*-values, *q*_*cl*_(*T*), is basically temperature-independent except damping coefficients, which increase linearly with temperature. In addition, *q*_*cl*_ generally does not change sign and is always real and positive. From the results in Figure 7, we notice that for TB, the two modes show similar *T*-dependence and, no temperature-hysteresis was observed both for tubulins or microtubules for all temperatures. While temperature hysteresis is a manifestation of classical behavior trough structural transitions and conformational memory leading to kinetically trapped states. However, the absence of such hysteresis is compatible with both quantum and classical regime. Future experiments with isotopic substitution (H_2_O → D_2_O) and eventual power laser dependence can definitively resolve the quantum/classical origin of the observed asymmetry. In fact, such type of experiments can discriminate between candidate mechanisms. Or, time-resolved Raman spectroscopy could determine whether phase-coherent transients accompany the steady-state asymmetry. Polarization-resolved measurements could isolate tensorial Raman matrix elements *M*_*d*_ and *M*_*c*_. Isotopic substitution, for example Deuterium substitution H_2_O → D_2_O [38-40], or selective mutagenesis could perturb specific vibrational coordinates entering in *H*_0_. Measurements under altered hydration, ionic strength, or nucleotide loading could directly test whether the continuum parameters *V* and *ρ* are environmentally controlled. On the theoretical side, atomistic normal-mode and open-quantum-system calculations could convert the phenomenological parameters extracted here into microscopic couplings.

Overall, the data support the view that TBs and MTs are not merely passive biomolecular scatterers, but structured dissipative media whose vibrational spectra encode interference between internal modes and environmental continua. The Fano framework therefore provides both a quantitative language for interpreting the present Raman results and a broader platform for investigating dynamical organization in cytoskeletal matter.

However, it is important to note that the experimentally supported interpretation remains phenomenological: Fano fitting captures spectral asymmetry well, but does not uniquely determine the microscopic origin of the continuum states. While Fano resonance provides an attractive framework for describing asymmetric Raman peaks, alternative explanations must be considered: i) Peak overlap from unresolved vibrational substructures, especially in densely packed protein environments; ii) Hydration-dependent spectral broadening, which introduces non-Lorentzian line shapes without requiring quantum interference; iii) Conformational heterogeneity, where multiple protein substrates contribute slightly shifted vibrational signatures.

We hope, current experimental work should stimulate new additional tasks in such directions.

## V. CONCLUSIONS

The present investigation demonstrated that Raman spectra of tubulin and microtubules contain reproducible asymmetric line shapes up to physiological temperatures.

These asymmetries can be quantitatively modeled using Fano resonance formalism, and the most plausible physical origin is the coupling between localized vibrational modes and broader electronic or environmental continua.

Within this derivation, the persistence of finite *q* values up to physiological temperature implies that the ratio of coherent resonant Raman scattering to dissipative continuum scattering remains non-negligible. This supports the view that MTs and TBs behave as structured open vibrational systems rather than featureless overdamped biomolecular media, denoting possible intriguing mechanisms of exploiting noisy biological environments.

In turn, since in biological systems, classical structural heterogeneity and solvent effects may contribute significantly, futher dedicated investigations could distinguish purely quantum interference from complex classical line-shape effects, since this question in Fano resonance remains an open methodological challenge.

## ACKNOWLEDGMENTS

The author wishes to thank Horiba Italy srl, M. Gaggini (CNR-IFC) and L. Sabatino (CNR-IFC) and F. Brandi (CNR-INO) for useful support.

The study was entirely carried out within the PNRR MUR project PE0000023-NQSTI financed by the European Union–Next Generation EU.

## APPENDIX

We consider the model as given by Fano-type model with dissipation, where the total Hamiltonian [41-42] *H = H*_0_ + *H*_*c*_ + *H*_int_is as follows:

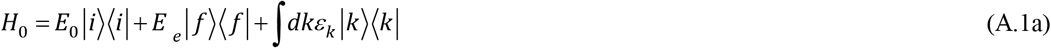

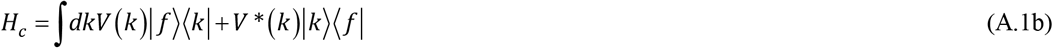

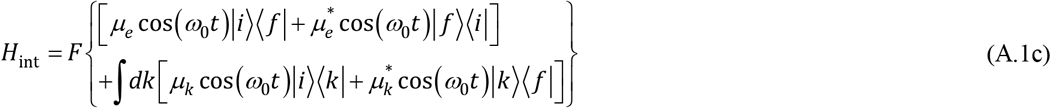

where *H*_0_ is the site Hamiltonian, *H*_c_ is the coupling of the excited state to the continuum and *H*_int_ is the interaction with the incident radiation field. A Liouvillian spectral decomposition could provide a deeper insight into the dynamics of dechorence and the temperature-dependent emergence of Fano lineshapes revealing how the asymmetry parameter *q*(*T*) and linewidth Γ(*T*) emerge from the eigenvalue spectrum of the Liouvillian superoperator. For this purpose it may be useful to handle the Fano-like Hamiltonian A.1, through a Feshbach-Fano procedure, keeping in mind that a Markovian expansion generates an effective non-Hermitian Hamiltonian [2, 43-45]:

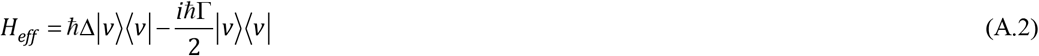

where Δ=*ω*_*L*_-*ω*_0_ include a shift term connected to coupling constant *V*. By projecting out the continuum degrees of freedom using the standard Feshbach-Fano procedure, the discrete vibrational mode acquires a complex self-energy Σ=Δ*-i*Γ/2. The resulting effective non-Hermitian Hamiltonian *H*_*eff*_ governs the reduced dynamics and forms the basis for the Liouvillian spectral description discussed below. The relaxation and dephasing processes consists of a Liouvillian equation describing the dynamics of the systems as

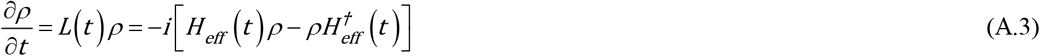

Eq. (A.3) generates a matrix which depends on the basis choice, which most important eigenvalue is *λ= −γ* +*i*ω_*eff*,_ and Re(*λ*)=−*γ*=−2*πV*^2^*ρ* is always negative.

To connect the phenomenological asymmetry parameter *q*(*T*) directly to the Hamiltonian (A.1), we derive the Raman scattering amplitude starting from the standard Kramers–Heisenberg formalism. This establishes how the experimentally fitted Fano line shape emerges from interference between discrete vibrational excitations and continuum environmental states in tubulin and microtubules. For incident radiation of frequency *ω*_*L*_ and scattered radiation of frequency *ω*_*S*_, the second-order Raman transition amplitude from an initial state ∣*i*⟩ to a final state ∣*f*⟩ is

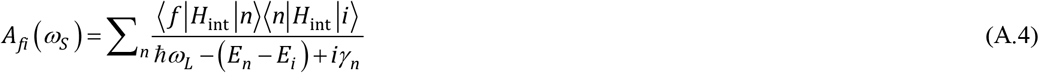

where the sum runs over intermediate virtual states ∣*n*⟩, and *γ*_*n*_ represents their damping width. In the present system, the intermediate manifold contains both a discrete vibrationally active channel and a quasi-continuum of environmental states.

The intensity of the Raman signal is proportional to

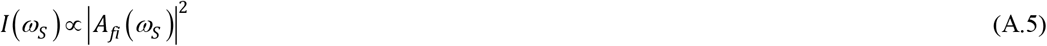

Now, we can decompose the total amplitude into two interfering contributions *A*(*ω* ) *= A*_*d*_ (*ω* ) + *A*_*c*_ (*ω* ) where *A*_*d*_(*ω*) denotes resonant discrete vibrational channel and *A*_*c*_(*ω*) the background continuum channel, respectively. The discrete contribution associated with a vibrational mode of frequency *ω*_*v*_ is

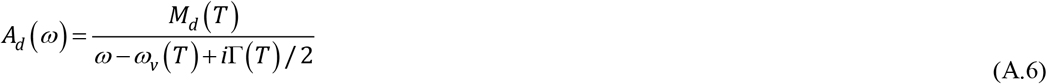

where *M*_*d*_(*T*) is the effective Raman matrix element of the discrete modes. The continuum contribution is slowly varying across the resonance and may be approximated as *A*_c_(*ω*)≈*M*_c_(*T*), where *M*_c_(*T*) is the non-resonant background scattering amplitude.

The role of coupling term in Fano-type Hamiltonian can be evidenced by the discrete–continuum interaction *H*_*c*_, Eq. (A.1). This term hybridizes the discrete vibration with continuum bath states. As in standard resonance theory, the self-energy correction is

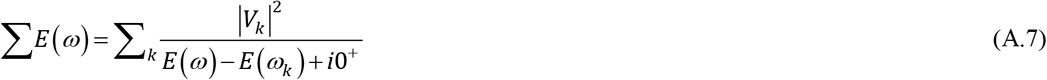

whose imaginary part gives the continuum-induced linewidth

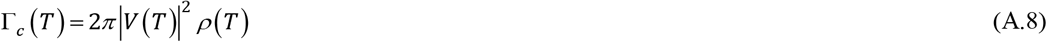

where *ρ*(*T*) is the effective density of continuum states at resonance. Thus the total linewidth includes not only continuum-induced contribution, but also intrinsic damping and pure dephasing. Under such picture we obtain Eq. (5) describing the asymmetry parameter is determined by the ratio of the discrete resonant amplitude to the continuum background amplitude, where *M*_*d*_(*T*): Raman matrix element of the discrete vibrational mode, *Mc*(*T*): continuum Raman scattering amplitude, *V*(*T*): coupling between discrete mode and continuum and *ρ*(*T*) the effective continuum density of states. Eq. (5) explicitly connects the fitted Fano parameter to the Hamiltonian terms: *M*_*d*_ and *M*_*c*_ originate from *H*_int_; *V* originates from *H*_c_; *ρ* is determined by the spectrum of *H*_0_.

In protein systems, all quantities entering *q*(*T*) may vary with temperature. Schematically *M*_*d*_(*T*) changes through modifications of polarizability derivatives, Eq. (2), and conformational flexibility. *V*(*T*) changes through altered coupling between local vibrations and environmental modes. *ρ*(*T*) changes through thermal population and restructuring of hydration-shell and collective lattice modes. *M*_*c*_(*T*) changes through background scattering from fluctuating protein and solvent coordinates. Therefore, the measured temperature dependence of the Fano asymmetry provides direct information on how biological self-assembly reorganizes vibrational coupling pathways. Consequently, shifts in the observed *q* values and linewidths are expected signatures of mesoscale structural organization.

